# tiRNA-Val promotes angiogenesis via Sirt1–Hif-1α axis in mice with diabetic retinopathy

**DOI:** 10.1101/2021.09.29.462300

**Authors:** Yan Xu, Haidong Zou, Qi Ding, Yuelan Zou, Chun Tang, Yuyu Lu, Xun Xu

**Author notes:** Corresponding author: Name:Xun Xu.

## Abstract

Diabetic retinopathy (DR) is a specific microvascular complication arising from diabetes, and its pathogenesis is notcompletely understood. tRNA-derived stress-induced RNAs (tiRNAs), a new type of small noncoding RNA generated by specific cleavage of tRNAs, has become a promising target forseveral diseases. However, the regulatory function of tiRNAs in DR and its detailed mechanism remain unknown. Here, we analyzed the tiRNA profiles of normal and DR retinal tissues. The expression level of tiRNA-Val was significantly upregulated in DR retinal tissues. Consistently, tiRNA-Val was upregulated in human retinal microvascular endothelial cells (HRMECs) under high glucose conditions. The overexpression of tiRNA-Val enhanced cell proliferation and inhibited cell apoptosis in HRMECs, but the knockdown of tiRNA-Val decreased cell proliferation and promoted cell apoptosis. Mechanistically, tiRNA-Val, derived from mature tRNA-Val with Ang cleavage, decreased Sirt1 expression level by interacting with sirt1 3’UTR, leading to the accumulation of Hif-1α, a key target for DR. In addition, subretinal injection of adeno-associated virus to knock down tiRNA-Val in DR mice ameliorated the symptoms of DR. Therefore, these data suggest that tiRNA-Val is a potential target in treating diabetic retinopathy.

## Introduction

Diabetic retinopathy (DR) is a common and a specific microvascular complication of diabetes(1), and it remains the leading cause of preventable blindness in working-aged people(2). It has been reported that one-third of those people with diabetes have an increased risk of life-threatening systemic vascular complications, such as stroke, coronary heart disease, and heart failure(3, 4). However, the pathogenesis of the onset of DR disease is notcompletely understoodas of yet.

Noncoding RNAs (ncRNAs) have emerged as critical regulators of various biological processes in DR, such as cell proliferation, cell motility,immune and inflammatory responses(5). For example, the expression of MIAT, a long noncoding RNA (lncRNA), increased in diabetic retinas, while MIAT knockdown ameliorated diabetes mellitus-induced retinal microvascular dysfunction(6). miRNA-138-5p is expressed at low levels in the retinal tissues of DR rats and it regulates early DR by promoting cell proliferation by targeting NOVA1(7). Recently, tRNA cleavage products have been identified asfunctional noncoding RNAs, called tRNA-derived stress-induced RNAs (tiRNAs), tRNA-derived RNA fragments (tRFs), or tRNA-derived small RNAs (tsRNAs)(8–10). tiRNAs are generated by specific cleavage in the anticodon loops of mature tRNAs or pre-tRNAs and are 31-40 bases long(11). The expression pattern of tiRNAs does not correspond to cognate tRNA levels, demonstrating that tiRNAs are not degradation products and precisely regulate noncoding RNAs(12). tiRNAs are an emerging class of regulatory non-coding RNAs that play important roles in regulating a variety of biological processes, such as competition for ribosomes(13), destabilizing YBX1-Bound mRNAs(14), and target mRNAs(15). However, the role of tiRNAs in DR is yet to be elucidated.

In this study, we constructed a DR mouse model with STZ-induced diabetes to analyze tiRNA profile of normal and DR retinal tissues. The expression level of tiRNA-Val was significantly upregulated in DR retinal tissues and in human retinal microvascular endothelial cells (HRMECs) under high glucose condition. tiRNA-Val enhanced cell proliferation and inhibited apoptosis in HRMECs. In addition, tiRNA-Val, derived from mature tRNA-Val, decreased Sirt1 expression level by interacting with Sirt1 3’UTR, leading to the accumulation of Hif-1α. Moreover, the knockdown of tiRNA-Val in retinal tissuesdrastically ameliorated the symptoms of DR *in vivo*. tiRNA-Val gene may be a potential target for diabetic retinopathy.

## Methods

### Cell lines and cell culture

HRMECs were purchased from American type culture collection (ATCC). HRMECs were cultured in Dulbecco’s modified eagle’s medium (DMEM) (Sigma-Aldrich, USA) supplemented with 1% penicillin/streptomycin (100 mg/L, Gibco, USA) and 10% heat-inactivated fetal bovine serum (FBS) (Gibco, USA) at 37°C in 5% CO_2_ atmosphere. For normal glucose and high glucose conditions, 5 mMand 33 mM D-glucose (Gibco, USA) were added to the medium for 48 h, respectively.

### Animals

All animal experiments were approved by the Institutional Animal Care and Use Committee of Shanghai General Hospital and were performed in accordance with the ARVO Statement for the Use of Animals in Ophthalmic and Vision Research. C57BL/6 male mice were purchased from Shanghai Model Organisms Center. The animals were housed in cages with free access to regular diet and water in a room at 22±1°C on a 12 h light/dark cycle. When the mice reached 20–25 g body weight (~2 months of age), they were randomlyassigned into diabetic or nondiabetic group. Diabetes was induced by five sequential daily intraperitoneal injections of a freshly prepared solution of streptozotocin in citrate buffer (pH 4.5) at 45 mg/kg body weight. Mice with random blood glucose levels ≥16.7 mmol/L at 2 weeks post-STZ were assigned to the diabetes group and the diabetes duration commenced. The animals had free access to food and water. Retinal tissues were harvested at 9 months of diabetes for protein extraction, RNA extraction, and retinal histopathology. Fasting blood glucose levels were determined repeatedly prior to the 3-month assessment.

For subretinal injection, adeno-associated virus (AAV) vector containing sh-tiRNA-Val under the control of chimeric CMV/chicken β-actin promoter was constructed. The vectors were administered viasubretinal injection two weeks before STZ induction of diabetes. C57BL/6 male mice were anesthetized and subretinally injected with 1μL solution containing 10^11^ particles ofsh-tiRNA-Val AAV, as previously described(16). The solution was injected onlyin one eye for each animal, while the contralateral eye was used as a control. Retinal tissues were harvested after 9 months of diabetes.

### Cell transfection

tiRNA-Val mimics, tiRNA-Val inhibitors, and corresponding negative controls were purchased fromSangon Biotech (Shanghai, China). Lipofectamine 3000 transfection reagent (Invitrogen, USA) was used for cell transfection according to the manufacturer’s instructions. The final concentrations of tiRNA-Val mimics andtiRNA-Val inhibitors were 50 nM, respectively.

### Cell proliferation assay

Cell viability was assessed using CCK-8 assay (Cell Counting Kit-8, Sigma-Aldrich, USA) according to the manufacturer’s instructions. Briefly, 5×10^3^ cells/well were seeded into 96-well plates. Proliferative activity was determined at the end of different experimental periods (24 h, 48 h, 72 h, and 96 h). When the medium changed from red to yellow, the absorbance value at a wavelength of 450 nm was detected using an enzyme-linked immunosorbent assay reader (Thermo Fisher Scientific, USA). The experiment was performed at least three times with similar results.

### Transwell migration assay

The migratory ability of HRMECs was assessed using 24-well transwell migration chambers (8 μm size, Corning, USA). Briefly, 5×10^4^ cells/well were resuspended in 200 μL serum-free DMEM and inoculated evenly into the inner chambers. The bottom chambers were replenished with 500 μLof DMEM containing 20% FBS as an attractant. After 24 h, the cells migrated to the lower chamber through the hole, fixed with 4% paraformaldehyde, and then stained with 0.1% crystal violet.

### Western blotting

Cell lysates or mouse tissues were prepared using 1× cell lysis buffer (Cell Signaling Technology, USA) with 1 mM phenylmethylsulfonyl fluoride (PMSF; Sigma-Aldrich, USA). Protein lysate of 10-20 μg was run on 10–15% SDS-PAGE gel and transferred toa PVDF membrane (Roche, USA). The membrane was incubated for 60 min at room temperature in 5% BSA solution. The following antibodies were used for the detection of protein expression: actin (1:1,000,Sigma, USA), angiogenin (Ang) (1:1,000,Abcam, USA), VEGF (1:1,000,Thermo Fisher Scientific, USA),ZO-1 (1:1,000,Thermo Fisher Scientific, USA), ICAM-1 (1:1,000,Abcam, USA), Sirt1 (1:1,000,Cell Signaling Technology, USA), and Hif-1α (1:1,000,Cell Signaling Technology, USA). Anti-rabbit and anti-mouse peroxidase-conjugated secondary antibodies (1:2,000,Cell Signaling Technology, USA) were purchased from Jackson Immunoresearch, and the signal was visualized using western blotting luminol reagent (Thermo Fisher Scientific, USA).

### Quantification of mRNA by RT-qPCR

Total RNA was isolated from cultured cells or mouse tissues using TRIzol reagent (Thermo Fisher Scientific, USA) according to the manufacturer’s instructions. For mRNA quantification, cDNA was synthesized using SuperScript IV Reverse Transcriptase (Thermo Fisher Scientific, USA) with random primers. RT-qPCR was performed using SYBR Green method. The primers used for amplification are listed in Supporting Information Table S1, and each experiment was repeated at least three times independently. The mRNA expression levels were calculated using β-actin as an internal control.

### Quantification of tiRNA by TaqMan RT-qPCR

TaqMan RT-qPCR for specific quantification of tiRNA was performed as previously described. Briefly, total RNA was treated with T4 PNK (New England Biolabs, UK), followed by ligation to 3’-RNA adapter using T4 RNA ligase. Ligated RNA was then subjected to TaqMan RT-qPCR using SuperScript IV Reverse Transcriptase, 200 nM of TaqMan probe targeting the boundary of target RNA and 3’-adapter, and specific forward and reverse primers. The expression of tiRNA was calculated using 5S RNA as an internal control. The sequences of the TaqMan probes and primers are listed in Table S2 of the Supporting Information.

### RNA cleavage reaction *in vitro*

RNA cleavage was performed as previously described(17). Briefly,the incubation mixtures contained 20 μg of total RNA extracted from HRMEC, 30 mM HEPES, pH 6.8, 30 mM NaCl, 0.001% BSA, and recombinant human angiogenin protein (R&D Systems, USA) at concentrations of 0.1μM, 0.2μM, 0.5μM, 1.0μM, and 2.0 μM. Incubation was performed at 37°C for 30 min. The cleaved products were recovered through phenol-chloroform extraction and ethanol precipitation. Then, the products were analyzed through northern blotting.

### Northern blotting

Northern blotting for specific detection of small RNA was performed as previously described(18). Briefly, total RNA was separated using 15% urea PAGE. Gels were stained with SYBR Gold nucleic acid gel stain (Thermo Fisher Scientific, USA) and immediately imaged and transferred to positively charged nylon membranes (Roche, Switzerland). Subsequently, the membranes were air-dried and UV-crosslinked. The membranes were pre-hybridized with DIG Easy Hyb buffer (Roche, Switzerland) for at least 1 h at 45°C. For the detection of specific small RNAs, the membranes were incubated overnight (12-16 h) at 45°C with 10 nM 3’-DIG-labeled oligonucleotide probes synthesized by Sangon Biotech (Shanghai, China), as shown in Supporting Information Table S3. The membranes were washed twice with low stringent buffer (2× SSC with 0.1% (w/v) SDS) at 37°C for 15 min each, then rinsed twice with high stringent buffer (0.1× SSC with 0.1% (w/v) SDS) at 37°C for 5 min each, and finally rinsed in washing buffer (1× SSC) for 10 min. Following the washes, the membranes were transferred onto 1× blocking buffer (Roche) and incubated at room temperature for 2-3 h, after which DIG antibody (Roche) was added to the blocking buffer at a ratio of 1:10,000 and incubated for an additional 1/2 min at room temperature. The membranes were then washed four times in DIG washing buffer (1 × maleic acid buffer, 0.3% Tween-20) for 15 min each, rinsed in DIG detection buffer (0.1 M Tris-HCl, 0.1 M NaCl, pH 9.5) for 5 min, and then coated with CSPD ready-to-use reagent (Roche, Switzerland). The membranes were incubated in the dark with CSPD reagent for 15 min at 37°C before imaging using the Carestream imaging system.

### Luciferase assay

HEK293T cells in a 24-well plate were cotransfected with pSIF-GFP or the indicated plasmids expressing tiRNA (0.8 μg/well), pRL-Sirt1-3’ UTR (pRL-TK vector containing Sirt1 3’UTR) or pRLSirt1-3’UTRm (pRL-TK vector containing mutant Sirt1 3’UTR) (0.1 μg/well), and pSV40-β-gal (Promega, Madison, WI, USA) (0.1 μg/well) using lipofectamine 3000. HERMEC cells in a 24-well plate were co-transfected with the indicated tiRNA mimics, pRL-Sirt1-3’UTR (0.1 μg/well), and pSV40-β-gal (0.1 μg/well) using lipofectamine 3000. After transfection for 72 h, the cells were harvested for luciferase assay as previously described(17).

### Statistical Analysis

Quantitative data are represented as mean ±SD. All images are representative of the studies with three to nine animals per group. Paired Student’s *t*-test was used to assess the significant difference between the two groups. Statistical significance was set at *p* ≤ 0.05.

## Results

### tiRNA profile in DR retinal tissues from mice

We constructed a mouse model with diabetic retinopathyaccording to a previously described method (19). The average fasting blood glucose level in DR mice was 19.0 mmol/L, which is far higher than that in normal mice (4.8 mmol/L) (Fig 1a). The mRNA expression levels of VEGF and ICAM-1 were significantly upregulated in DR retinal tissues, while the mRNA expression level of ZO-1 was significantly downregulated (Fig 1b). The protein levels of VEGF, ICAM-1, and ZO-1 also changed based on the mRNA level (Fig 1c). To evaluate the degeneration of retinal neurons, we examined retinal ganglion cell layer (GCL) after 9 months of diabetes. Diabetic mice experienced 10% loss of neurons in retinal GCL compared to that in non-diabetic mice (Fig 1d).

**Fig. 1.**
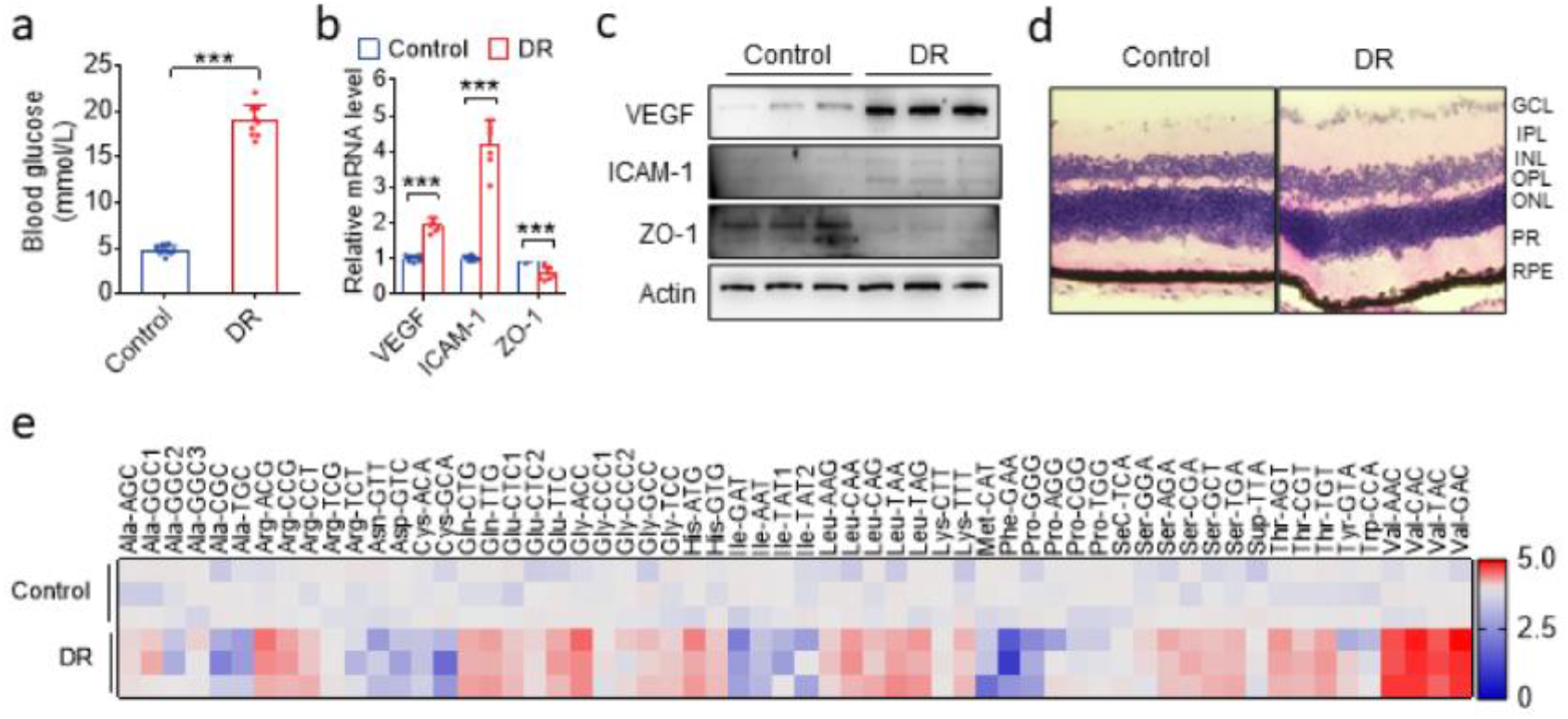
tiRNA profile between normal and DR retinal tissues in mice. (a) Blood glucose level in normal and DR mice.Data are represented as the mean ± SD, n = 9, \*\*\**p*< 0.001 *vs*. normal group.Statistical significance was assessed by two-tailed Student’s *t*-test. (b)qRT-PCR analysis of *VEGF*, *ICAM-1*, and *ZO-1 levels* in the entire retina of DR mice. Data are represented as the mean ± SD, n = 6, \*\*\**p*< 0.001 *vs*. normal group. Statistical significance was assessed by two-tailed Student’s *t*-test. (c)Western blotting analysis of *VEGF*, *ICAM-1*, and *ZO-1* expressionin the entire retina of normal and DR mice. (d)Representative micrographs of H&E staining of theretinatissue in mice treated as indicated. (e) Heatmap of differently expressed tiRNAs between normal and DR mice retinal tissues by TaqMan RT-qPCR. GCL: ganglion cell layer; IPL:inner plexiform layer; INL:inner nuclear layer;OPL:outer plexiform layer;ONL:outer nuclear layer;PR:photoreceptors; RPE:retinal pigment epithelium;DR: diabetic retinopathy.

To explore the physiological relevance of tiRNAs, TaqMan RT-qPCR quantification of all the tiRNAs that cleaved at the anticodon loop was performed for DR retinal tissues of mice. As shown in Fig 1e, the tiRNA profile was significantly altered in the retinal tissues of DR mice, especially tiRNA-Val, which was markedly upregulated. Therefore, we chose tiRNA-Val as a candidate for this study.

### tiRNA-Val was upregulated in DR retinal tissues and HRMEC at high glucose condition

tiRNA-Val was derived from mature tRNA-Val, which was cleaved at the anticodon loop with RNase (Fig 2a). We analyzed the expression level of tiRNA-Val through northern blotting. As shown in figure 2b, no significant differences were observed in the expression level of mature total tRNA-Val, but tiRNA-Val was significantly upregulated in the retinal tissues of DR mice. The mRNA and protein levels of VEGF, ICAM-1, and ZO-1 significantly changed in theretinal tissues of DR mouse (Fig 2c and 2d). Furthermore, HRMECs treated with the indicated concentrations of glucose were used to simulate various diabetic conditions(20). HRMECs were cultured under normal glucose (5 mM) or high-glucose (33 mM) conditions. The expression level of tiRNA-Val was upregulated in HRMECs under high glucose conditions (Fig 2e).

**Fig. 2.**
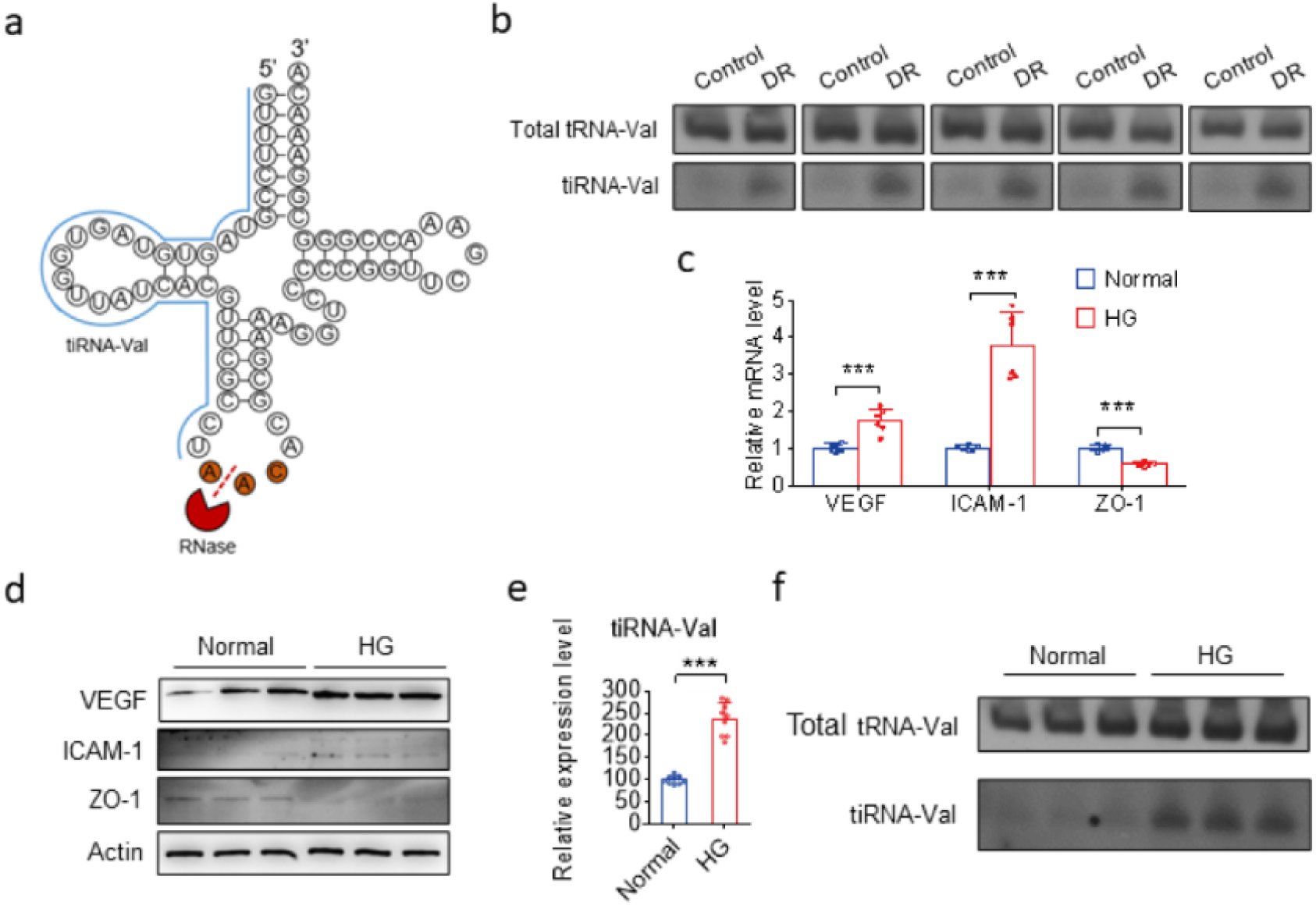
The expression level of tiRNA-Val was upregulated in DR mice and high glucose cell model. (a)Structure of tiRNA-Val and total tRNA-Val. (b) Expression level of tiRNA-Val identified in five pairs of normal and DR retinal tissues by northern blot. (c)qRT-PCR analysis of *VEGF*, *ICAM-1*, and *ZO-1* levels in normal and high glucose HRMEC.Data are represented as the mean ± SD, n = 6, *** *p*< 0.001 *vs*. normal group. Statistical significance was assessed by two-tailed Student’s Atest. (d) Western blotting analysis of *VEGF*, *ICAM-1*, and *ZO-1* expression in HRMECs treated mice as indicated. (e)Expression level of tiRNA-Val identified by TaqMan RT-qPCR in normal and high glucose HRMEC. Data are represented as the mean ± SD, n = 9, \*\*\**p*< 0.001 *vs*. normal group. Statistical significance was assessed by two-tailed Student’s Atest. (f)Expression level of tiRNA-Val identified in 3 pairs of normal and high glucose HRMEC by northern blotting.DR: diabetic retinopathy; NG: normal glucose;HG:high glucose.

### tiRNA-Val enhance cell proliferation in HRMEC

DR is a proliferative manifestation of the retinaaccompanied by the growth of abnormal new blood vessels(21). We investigated the regulatory function of tiRNA-Val in cell proliferation by performing tiRNA-Val transfection (Fig. 3a). In view of the high expression of tiRNA-Val in the retinal tissues of DR mice and high glucose cell models, we examined the effect of tiRNA-Val on proliferation and migration of HRMECs.tiRNA-Val mimics and tiRNA-Val inhibitors were transfected into HRMECs, respectively, followed by CCK-8 and transwell migration assays. As shown in Fig. 3b, the viability of HRMECs increased markedly by transfection with tiRNA-Val mimics, and the enhanced effect of tiRNA-Val mimics on cell proliferation was observed beginning at 48 h. HRMECs transfected with tiRNA-Val mimics migrated significantly faster than those in the cells transfected with the negative control (Fig. 3c). To further investigate the effect of tiRNA-Val on cell apoptosis in HRMECs, FITC Annexin V apoptosis detection was performed. It was found that cells transfected with tiRNA-Val mimics could significantly inhibit cell apoptosis compared to that in cells transfected with the negative control (Fig. 3d). In addition, HRMECs transfected with tiRNA-Val inhibitors to knock down tiRNA-Val decreased cell proliferation and migration (Fig. 3e-g), but promoted cell apoptosis (Fig. 3h).

**Fig. 3.**
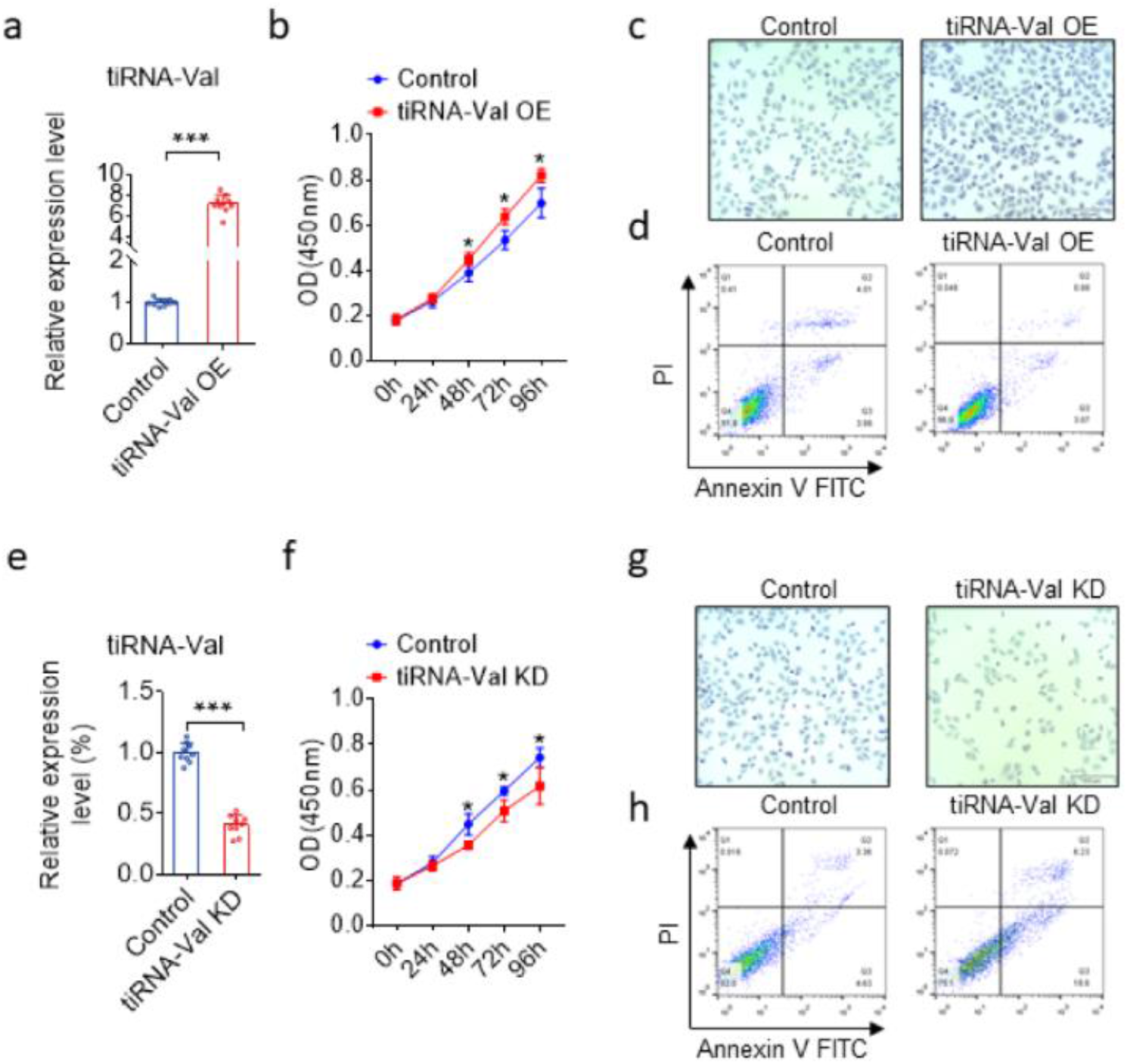
The regulatory function of tiRNA-Val in HRMEC cells. (a) TaqMan RT-qPCR analysis of tiRNA-Val expression in HRMEC cells transfected with tiRNA-Val mimics and scramble sequence RNA. HRMEC cells transfected withscramble sequence RNA as control group. Data are represented as the mean ± SD, n = 10, \*\*\**p*< 0.001 *vs*. control group. Statistical significance was assessed by two-tailed Student’s *t*-test. (b)CCK-8 assay for HRMEC cells transfected with tiRNA-Val mimics compared to scramble sequence RNA. HRMEC cells transfected withscramble sequence RNA as control group. Data are represented as the mean ± SD, n = 9, \*\*\**p*< 0.001 *vs*. control group. Statistical significance was assessed by two-tailed Student’s *t*-test. (c)Migration assay for HERMEC cells transfected with tiRNA-Val mimics and scramble sequence RNA. (d) Detection of apoptosis by concurrent staining with Annexin V-FITC and PI. HRMEC cells transfectedwithtiRNA-Val mimics (left panel) or scramble sequence RNA (right panel). Cells were subsequently stained with Annexin V-FITC conjugate and PI and were measured by flow cytometry. Live cells were both Annexin V and PI negative. At early stage of apoptosis, the cells bound Annexin V while still excluding PI. At the late stage of apoptosis, they bound Annexin V-FITC and stained brightly with PI. (e) TaqMan RT-qPCR analysis of tiRNA-Val expression in HRMEC cells transfected with siRNA of tiRNA-Val and scramble sequence RNA. HRMEC cells transfected withscramble sequence RNA as control group. Data are represented as the mean ± SD, n = 9, \*\*\**p*< 0.001 *vs*. control group. Statistical significance was assessed by two-tailed Student’s *t-* test.(f)CCK-8 assay for HRMEC cells transfected with siRNA of tiRNA-Val compared to scramble sequence RNA. (g)Migration assay for HRMEC cells transfected with siRNA of tiRNA-Val compared to scramble sequence RNA.(h) Detection of apoptosis by concurrent staining with Annexin V-FITC and PI. HRMEC cells transfectedwithsi-tiRNA-Val (left panel) or scramble sequence RNA (right panel). Cells were subsequently stained with Annexin V-FITC conjugate and PI as described in (b).

### Ang cleaves tRNA-Val to produce tiRNA-Val in mouse retinal tissues and HRMEC cell models

Previousstudies have shown that tiRNA production is dependent on Ang, which is the fifth member of the RNase A superfamily(17, 22). The mRNA and protein levels significantly increased in the retinal tissues of DR mice (Fig. 4a and 4b). To test whether Ang could cleave tRNA-Val, total RNA from HRMECs was incubated with recombinant Ang *in vitro*. Northern blotting results showed that intact tRNA-Val was cleaved into short tRNA fragments of the length of tiRNA-Val (Fig. 4c). To test whether Ang could cleave tRNAs in cultured mammalian cells, HRMECs were transiently transfected with a plasmid expressing angiogenin. Total cellular RNA was extracted after transfection for 48 h. tiRNA-Val significantly increased in Ang-overexpressing cells (Fig. 4d-4e). However, tiRNA-Val levels were not detected in HRMECs transfected with Ang siRNA (Fig. 4f-4g). These results suggest that Ang is possibly an endonuclease for producing tiRNA-Val *in vivo*.

**Fig. 4.**
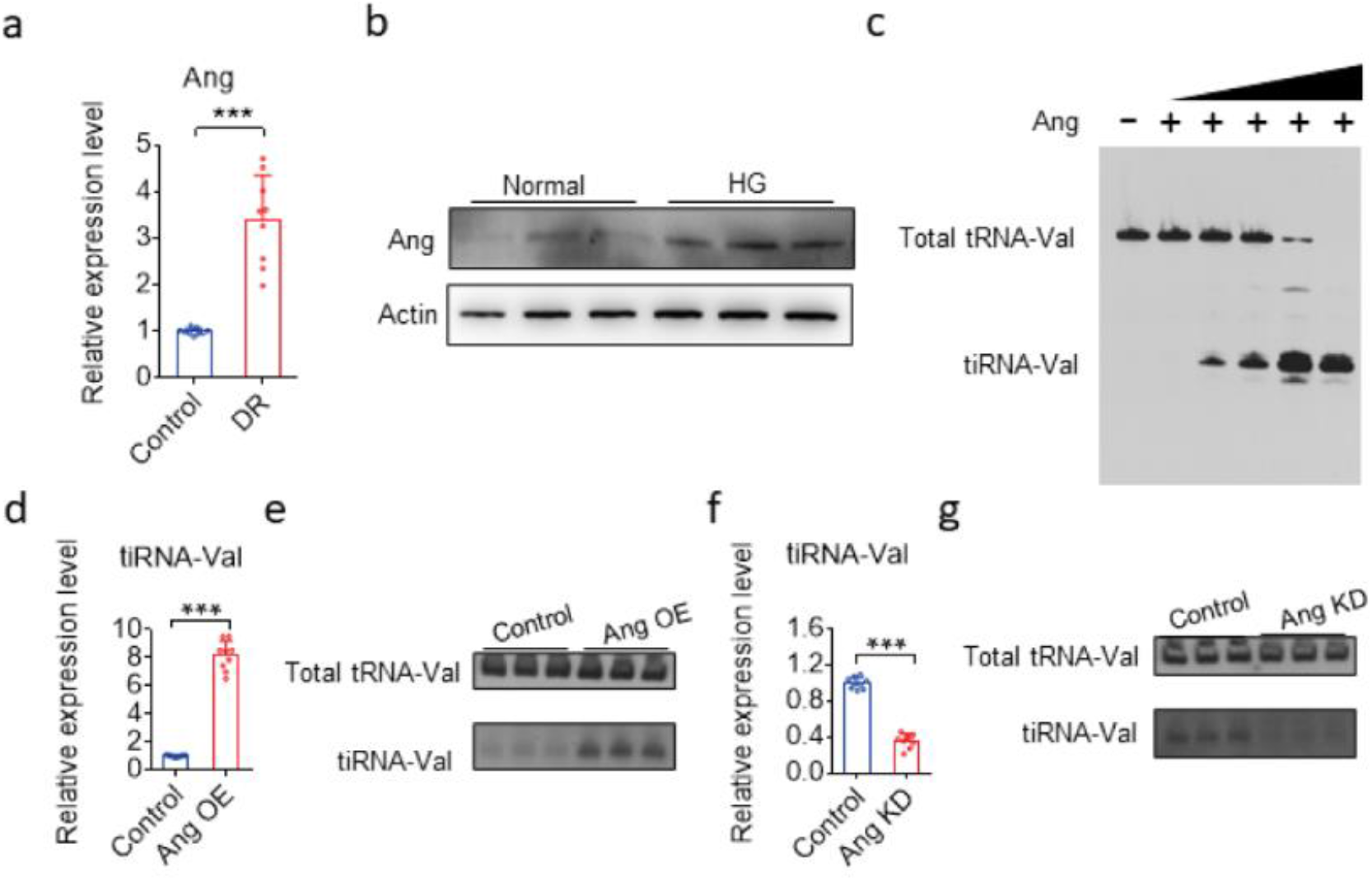
Ang cleaves tRNA-Val to produce tiRNA-Val in mice retinal tissues and HRMEC cell models. **(a)** TaqManqRT-PCR analysis of *angiogenin*(*Ang*)*levels* in the entire retina of DR mice. Data are represented as the mean ± SD, n = 9, \*\*\**p*< 0.001 *vs*. normal group. Statistical significance was assessed by two-tailed Student’s *t*-test. **(b)** Western blotting analysis of Ang expressionin the entire retina of normal and DR mice. **(c)**tiRNA-Val can be cleaved at the anticodon loop depending on the recombinant angiogenin(0.1, 0.2, 0.5, 1.0 and 2.0 μM) in vitro. **(d)**TaqMan RT-qPCR analysis of tiRNA-Val expression in HRMEC cells transfected with Ang overexpression plasmid and empty vector in normal glucose. HRMEC cells transfected withempty vector as control group. Data are represented as the mean ± SD, n = 9, \*\*\**p*< 0.001 *vs*. control group. Statistical significance was assessed by two-tailed Student’s *t*-test. **(e)**Overexpression of Ang in HRMEC with normal glucose to analyze tiRNA-Val level by northern blotting. **(f)** Knockdown of Ang in HRMEC with high glucose level to analyze tiRNA-Val level by northern blotting. **(g)**TaqMan RT-qPCR analysis of tiRNA-Val expression in HRMEC cells transfected with siRNA of Ang andscramble sequence RNA in high glucose. HRMEC cells transfected withscramble sequence RNA as control group. Data are represented as the mean ± SD, n = 9, \*\*\**p*< 0.001 *vs*. control group. Statistical significance was assessed by two-tailed Student’s *t*-test.

### tiRNA-Val increased Hif-1α expression level by interacting with Sirt1 3’UTR

tiRNAs are a new class of small RNAs with different mechanisms to regulate various cellular processes(12). Hif-1α is a key mediator and target of retinal neovascularization and diabetic retinopathy(23, 24), and during hypoxia, Sirtuin 1 (Sirt1) is downregulated, which allows the acetylation and activation of Hif-1α. We found that tiRNA-Val could pair with the 3’UTR of Sirt1 (Fig. 5a). Then, a luciferase reporter under the control of Sirt1 3’UTR was used to examine the effect of tiRNA-Val. As shown in Fig. 5b, the overexpression of tiRNA-Val significantly downregulated the activity of Sirt1 3’UTR, whereas the overexpression of tiRNA-Val had no effect on the mutant reporter. To further confirm whether tiRNA-Val targets Sirt1 3’UTR, a plasmid expressing mutant tiRNA-Val with ten mismatched bases was constructed, and we found that the mutant tiRNA-Val had no effect on the activity of Sirt1 3’UTR (Fig. 5c). To further examine the relationship among tiRNA-Val, Hif-1α, and Sirt1, we performed transfection with tiRNA-Val mimics and found that Hif-1α protein levels significantly increased, whereas Sirt1 protein levels decreased (Fig. 5d). Similarly, Hif-1α protein level was upregulated, but the protein level of Sirt1 significantly decreased in the retinal tissue of DR mice (Fig. 5e). These data demonstrate that tiRNA-Val decreased Sirt1 expression level by interacting with Sirt1 3’UTR leading to the accumulation of Hif-1α.

**Fig. 5.**
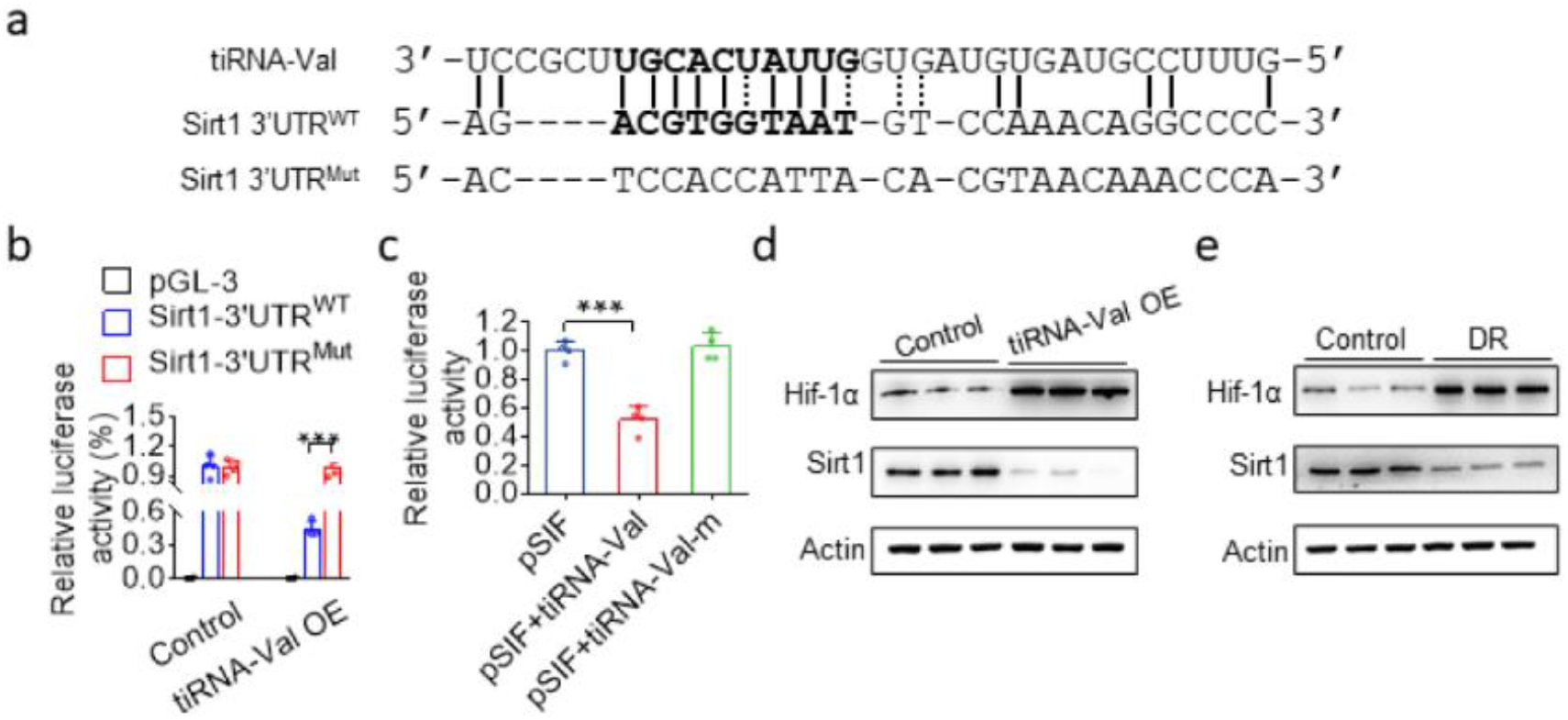
tiRNA-Val targets sirt1 3’UTR leading to the accumulation of Hif-1α. (a)Sequence alignment of tiRNA-Val with the 3’UTRs of Sirt1. The seed region of tiRNA-Val is indicated in bold.(b)Luciferase assay indicated the mutation of the predicted tiRNA-Val binding site in Sirt1 3’UTR,and it abrogated the repressive effect of tiRNA-Val on the activity of Sirt1 3’UTR. Data are represented as the mean ± SD, n = 4, \*\*\**p*< 0.001 *vs*. Sirt1 3’UTR^WT^ group. Statistical significance was assessed by two-tailed Student’s *t*-test. (c)Luciferase assay indicated the mutation of the tiRNA-Val seed region, and it abrogated the repressive effect of tiRNA-Val on the activity of Sirt1 3’UTR. Data are represented as the mean ± SD, n = 4, \*\*\**p*< 0.001 *vs. p*SIF+tiRNA-Val group. Statistical significance was assessed by two-tailed Student’s *t*-test. (d)Western blotting analysis of Hif-1 α and Sirt1 expression in HRMECs transfected withtiRNA-Val mimics and the scramble sequence RNA.(e)Western blotting analysis of Hif-1α and Sirt1 expression in the entire retina of normal and DR mice. WT: wile type; DR: diabetic retinopathy; Mut:mutation; OE: overexpression.

### Knockdown of tiRNA-Val ameliorates DR *in vivo*

To explore tiRNA*in vivo*, we knocked down tiRNA-Val in the subretinal space of DR mice with AAV -shtiRNA-Val. As shown in Fig. 6a and 6b, the expression level of tiRNA-Val decreased to 34.9%. The protein level of Sirt1 significantly increased when tiRNA-Val was knocked down, and Hif-1α was upregulated (Fig. 6c). Importantly, the mRNA and protein levels of VEGF and ICAM-1 were downregulated, while ZO-1 increased significantly (Fig. 6d and 6e). Moreover, the loss of neurons in GCL was recovered compared tothat in diabetic mice with control AAV (Fig 6f). These data demonstrated that the knockdown of tiRNA-Val ameliorated the symptoms of DR *in vivo*.

**Fig. 6.**
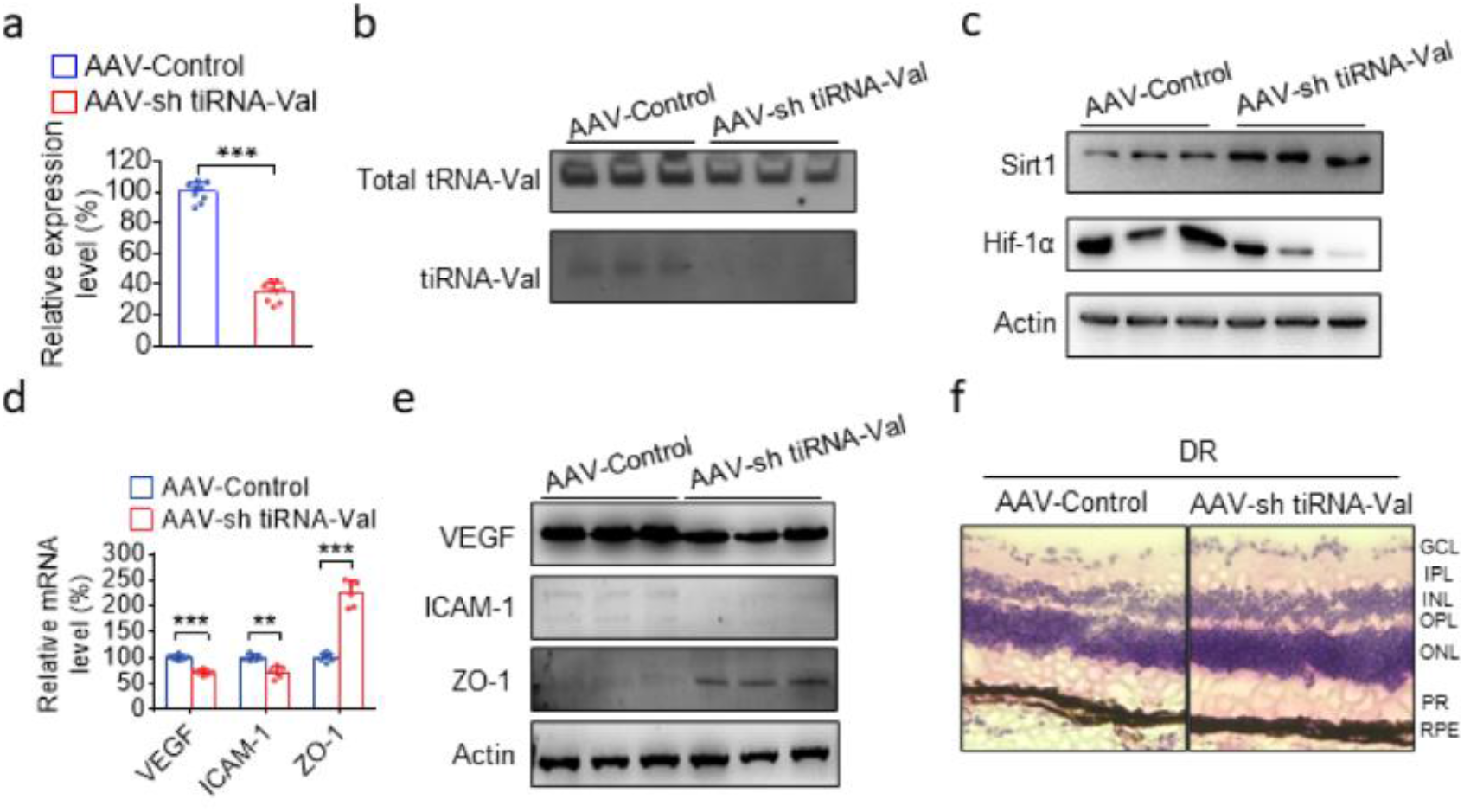
Knockdown of tiRNA-Val in subretinal space ameliorates thesymptoms of DR *in vivo*. (a)TaqManqRT-PCR analysis of tiRNA-Val levels in the entire retina of DR mice after shtiRNA-Val AAV injection. Data are represented as the mean ± SD, n = 9, \*\*\**p*< 0.001 *vs*. normal group. Statistical significance was assessed by two-tailed Student’s *t*-test. (b)Northern blot detection of tiRNA-Val level in DR mice retinal tissues after shtiRNA-Val AAV injection.(c)Western blotting analysis of Hif-1 α and Sirt1 expression in DR mice retinal tissues after shtiRNA-Val AAV injection. (d)TaqManqRT-PCR analysis *of VEGF*, *ICAM-1*, and *ZO-1* levels in the entire retina of DR mice after shtiRNA-Val AAV injection. Data are represented as the mean ± SD, n = 6, \*\*\**p*< 0.001 *vs*. normal group. Statistical significance was assessed by two-tailed Student’s *t*-test.(e)Western blotting analysis *of VEGF*, *ICAM-1*, and *ZO-l*expression in DR mice retinal tissues after shtiRNA-Val AAV injection. (f)Representative micrographs of H&E staining of retinatissue in DR mice after tiRNA-Val AAV injection. GCL: ganglion cell layer; IPL:inner plexiform layer; INL:inner nuclear layer;OPL:outer plexiform layer;ONL:outer nuclear layer;PR:photoreceptors; RPE:retinal pigment epithelium; DR: diabetic retinopathy.

## Discussion

In this study, we found that tRNA-derived small RNA, tiRNA-Val, was upregulated in the retinal tissues of DR mice. Ang, a member of the RNase A family, cleaved mature tRNA-Val to tiRNA-Val, which could enhance cell proliferation in HRMECs. Furthermore, we identified Sirt1 as the direct target of tiRNA-Val and demonstrated that tiRNA-Val negatively regulated Sirt1 in DR. It has been reported that hypoxia decreases Sirt1 expression, leading to the acetylation and activation of Hif-1α(25). Our findings showed that tiRNA-Val downregulated the expression level of Sirt1, leading to the accumulation of Hif-1α. The knockdown of tiRNA-Val in the subretinal space ameliorated DR via Sirt1-Hif-1α axis *in vivo* (Fig. 7). These results suggest that tiRNA-Val may represent a potential therapeutic target for the treatment of DR.

**Fig. 7.**
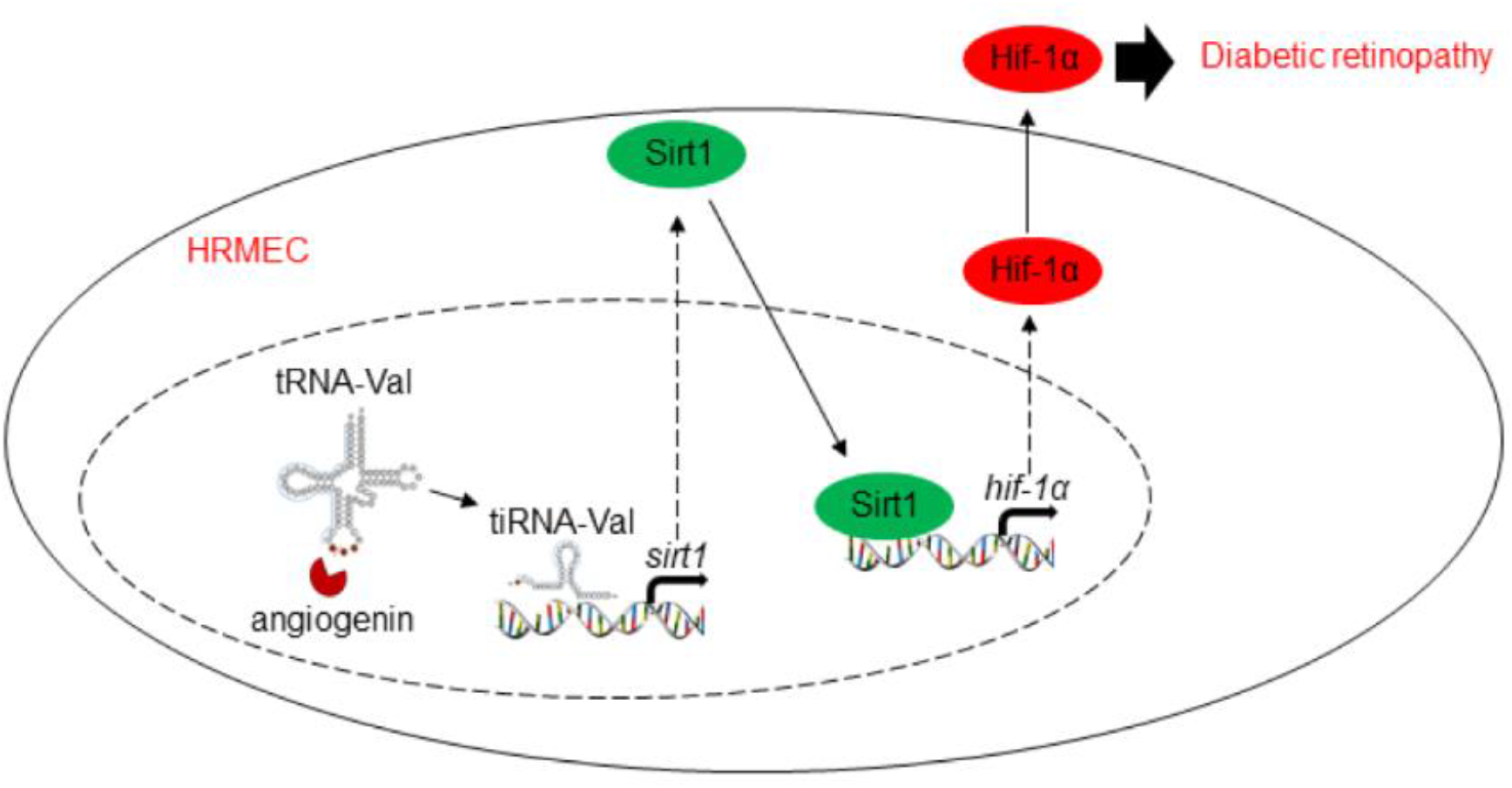
Model for tiRNA-Val regulating diabetic retinopathy via Sirt1-Hif-1α axis. Model for tiRNA-Val regulating diabetic retinopathy via Sirt1-Hif-1α axis

High-throughput sequencing has resulted in the discovery of a new class of small RNAs: tRFs and tiRNAs derived from tRNAs. tiRNAs are activated under stress conditions and they modulate the stress response(26). Although they are named stress fragments, they are detected under non-stressed conditions(27). tiRNAs span the entire evolutionary tree, and biological roles have been identified for some tiRNAs in subsets of organisms. For example, tiRNA-Ala can inhibit protein synthesis and promote stress granule formation in a phospho eIF2α independent manner, inhibiting translation by displacing the eukaryotic initiation factor eIF4G/A from mRNAs(22). A group of tiRNAs competitively bind to cytochrome c, protecting cells from apoptosis during osmotic stress cytochrome c(28). tiRNAs from the sperm contribute to intergenerational inheritance and alter the expression profile and RNA modifications of many genes(29). Here, we found that tiRNA-Val negatively regulated Sirt1 in DR by interacting withSirt1 3’UTR. Previous studies have shown that tRNA-derived fragments can repress endogenous genes to regulate cell proliferation and modulate DNA damage response(30, 31). It is possible that tiRNAs play a key role in regulating gene expression levels in miRNA pathway or take part in other mechanisms.

SIRT1 is a nicotinamide adenosine dinucleotide (NAD)-dependent multifunctional deacetylase that removes acetyl groups from many proteins that can be implicated in diabetes(32). It was reported that Sirt1 was downregulated in DR patients(33). Sirt1 regulated the expression of Hif-1α, especially under hypoxic condition; thus, it was involved in multiple biological processes associated with DR progression, such as apoptosis and proliferation(25). Here, we found that Sirt1 was downregulated by tiRNA-Val, leading to the accumulation of Hif-1α in HRMECs. Meanwhile, the knockdown of tiRNA-Val with shtiRNA-Val AAV subretinal injection ameliorates DR via Sirt1-Hif-1α axis *in vivo*.

In summary, we identified and characterized a small RNA, tiRNA-Val, that regulates diabetic retinopathy by modulating cell proliferation, and wehave shown a potential approach that can be used to improve diabetic retinopathy by knocking down tiRNA-Val.

## Abbreviations

DR: Diabetic retinopathy
tiRNAs: tRNA-derived stress-induced RNAs
HRMEC: Human retinal microvascular endothelial cell
ncRNAs: Noncoding RNAs

## Author contributions

YX and XX conceived and designed the study. YX and XX performed the experiments and drafted the manuscript. HZ, QD, YLZ, CT and YYL helped to analyze the data. XX supervised the experiments and revised the manuscript. All of the authors reviewed the manuscript and approved the final version.

## Compliance with ethical standards

